# Data proliferation, reconciliation, and synthesis in viral ecology

**DOI:** 10.1101/2021.01.14.426572

**Authors:** Rory Gibb, Gregory F. Albery, Daniel J. Becker, Liam Brierley, Ryan Connor, Tad A. Dallas, Evan A. Eskew, Maxwell J. Farrell, Angela L. Rasmussen, Sadie J. Ryan, Amy Sweeny, Colin J. Carlson, Timothée Poisot

## Abstract

The fields of viral ecology and evolution have rapidly expanded in the last two decades, driven by technological improvements, and motivated by efforts to discover potentially zoonotic wildlife viruses under the rubric of pandemic prevention. One consequence has been a massive proliferation of host-virus association data, which comprise the backbone of research in viral macroecology and zoonotic risk prediction. These data remain fragmented across numerous data portals and projects, each with their own scope, structure, and reporting standards. Here, we propose that synthesis of host-virus association data is a central challenge to improve our understanding of the global virome and develop foundational theory in viral ecology. To illustrate this, we build an open reconciled mammal-virus database from four key published datasets, applying a standardized taxonomy and metadata. We show that reconciling these datasets provides a substantially richer view of the mammal virome than that offered by any one individual database. We argue for a shift in best practice towards the incremental development and use of synthetic datasets in viral ecology research, both to improve comparability and replicability across studies, and to facilitate future efforts to use machine learning to predict the structure and dynamics of the global virome.

## Introduction

The emergence of SARS-CoV-2 was a harsh reminder that uncharacterized wildlife viruses can suddenly become globally relevant. Efforts to identify wildlife viruses with the potential to infect humans, and to predict spillover and emergence trajectories, are becoming more popular than ever (including with major scientific funders). However, the value of these efforts is limited by an incomplete understanding of the global virome (Wille et al. 2020). Significant knowledge gaps exist regarding the mechanisms of viral transmission and replication, host-pathogen associations and interactions, spillover pathways, and several other dimensions of viral emergence. Further, although billions of dollars have been invested in these scientific challenges over the last decade alone, much of the data relevant to these problems remains unsynthesized. Fragmented data access and a lack of standardization preclude an easy reconciliation process across data sources, making the whole less than the sum of its parts, and hindering synthetic research (Wyborn et al. 2018).

Here, we propose that data synthesis is a seminal challenge for translational work in viral ecology. This requires researchers to go beyond the usual steps of data collection and publication, to develop a community of practice that prioritizes data synthesis and reconciles semi-reproduced work across different teams and disciplines. As an illustrative example, we describe the analytical hurdles of working with **host-virus association data**, a format that characterizes the global virome as a bipartite network of hosts and viruses, with pairs connected by observed potential for infection. Recent studies highlight the central role for these data in efforts to understand viral macroecology and evolution (Carlson et al. 2019, Dallas et al. 2019, Albery et al. 2020), to predict zoonotic emergence risk (Han et al. 2015, 2016, Olival et al. 2017, Wardeh et al. 2020), and to anticipate the impacts of global environmental change on infectious disease (Carlson et al. 2020, Gibb et al. 2020, Johnson et al. 2020). Several bespoke datasets have been compiled to address these questions, and as interest in these topics has grown, so has the fragmentation of total knowledge across those datasets. To illustrate this problem (and a simple solution), we compare and reconcile four major host-virus association datasets, each of which is different enough that we anticipate the results of individual studies could be strongly shaped by choice of dataset.

### Four parts of one whole

Though host-pathogen association data exist in dozens of sources and repositories, there are at least four published datasets that each capture between 0.3% and 1.5% of the estimated 50,000 species of mammal viruses (Carlson et al. 2019). Differences among these datasets, especially with regards to available metadata and frequency of data updates, make them preferable for different purposes (Table 1), but may also complicate intercomparison and synthetic inference.

**Table 1.**
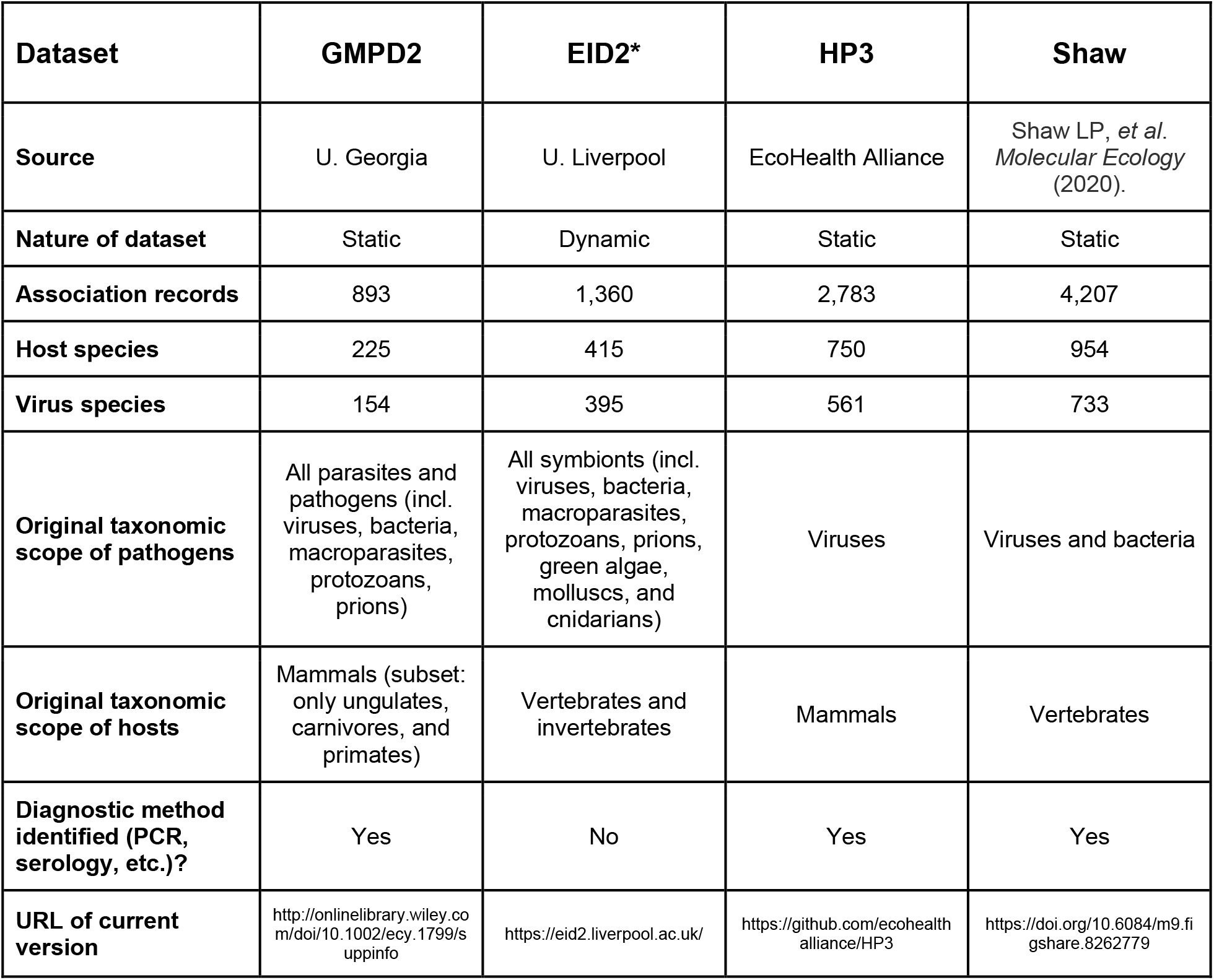
Available “big data” on host-virus associations, and major features of each dataset. Numbers of unique association records and host, virus, and pathogen species are all derived from the reconciled version presented in the CLOVER database, and therefore these numbers may differ from those presented in the main text (which are taken from the source data, or from self-reporting by the data curators). *Number of associations and taxa accurate as of 2015 static release in *Scientific Data* paper.

#### GMPD 2.0

The Global Mammal Parasite Database (Nunn and Altizer 2005), started in 1999 and now in its second public version (Stephens et al. 2017), emerged from continued efforts to compile mammal-parasite association data from published literature sources. Construction of the GMPD used a variety of similar strategies that combined host Latin names with a string of parasite-related terms to search online literature databases. Pertinent literature was then manually identified and relevant association and metadata compiled. The initial database was focused on primate hosts (Nunn and Altizer 2005), and expanded to include separate sections for ungulates (Ezenwa et al. 2006) and carnivores (Lindenfors et al. 2007). In 2017, GMPD 2.0 was released, which merged these three previously independent databases that were being independently maintained and updated (Stephens et al. 2017). The updated dataset encompasses 190 primate, 116 ungulate, and 158 carnivore species, and record their interactions with 2,412 unique “parasite” species, including 189 viruses, as well as bacteria, protozoa, helminths, arthropods, and fungi. Notable improvements in version 2 of the GMPD are the construction of a unified parasite taxonomy that bridges occurrence records across host taxa, the expansion of host-parasite association data along with georeferencing, and enhanced parasite trait data (e.g., transmission mode). The original data are available as a web resource (*www.mammalparasites.org*), and the data from GMPD version 2 can also be downloaded as static files from a data paper (Stephens et al. 2017). In addition, one subsection of the GMPD, named the “Global Primate Parasite Database,” has been independently maintained and regularly updated by Charles Nunn (data available at https://parasites.nunn-lab.org/). Consequently, the primate subsection of GMPD 2.0 includes papers published up to 2015, while the ungulate and carnivore subsections stop after 2010 (Stephens et al. 2017).

#### EID2

The ENHanCEd Infectious Diseases Database (EID2), curated by the University of Liverpool, may be the largest dynamic dataset of any symbiotic interactions (Wardeh et al. 2015). EID2 is compiled from automated, dynamic scrapes of two web sources: publication titles and abstracts indexed in the PubMed database and the NCBI Nucleotide Sequence database (along with its associated taxonomic metadata). The EID2 data is structured using the concepts of “carrier” and “cargo” rather than host and pathogen, as it includes a number of ecological interactions beyond the scope of normal host-pathogen interactions, including potentially unresolved mutualist or commensal associations. Interactions are stored as a geographic edgelist, where each carrier and cargo can also have locality information; additional metadata include the number of sequences in GenBank and related publications. EID2’s dynamic web interface (currently available through download on a limited query-by-query basis which researchers often manually bind or by personal correspondence with data curators) contains information encompassing 4,799 mammal “carrier” species and 70,614 microparasite or macroparasite “cargo” species, of which 9,605 are viruses (Wardeh et al. 2020). However, many researchers continue to use the static, open release of EID2 from a 2015 data paper (Wardeh et al. 2015), which we focus on here for comparative purposes as a stable version of the database available to the community of practice. The EID2 data were originally validated for completeness against GMPD 1.0.

#### HP3

The Host-Parasite Phylogeny Project dataset (HP3) was developed by EcoHealth Alliance over the better part of a decade. Published along with a landmark analysis of zoonotic spillover (Olival et al. 2017), the HP3 dataset consists of 2,805 associations between 754 mammal hosts and 586 virus species. These were compiled from literature published between 1940 and 2015, based on targeted searches of online reference databases. Complementary with the search strategy used for the GMPD, rather than starting with a list of host names, HP3 started with names of known mammal viruses listed in the International Committee on Taxonomy of Viruses (ICTV) database. These virus names along with their synonyms were then used as search terms to identify literature containing host-virus association data. To narrow search results for well-studied viruses, they included additional host range-related terms to identify relevant publications. Data collection and cleaning for HP3 began in 2010 and the database has been static since 2017; it can be obtained as a flat file in the published study’s data repository (Olival et al. 2017). HP3 includes a host-virus edgelist (see Glossary), separate files for host and virus taxonomy, and separate files for host and virus traits. Host-virus association records are provided with a note about method of identification (PCR, serology including specific methods, etc.), which may be useful for researchers interested in the different levels of confidence ascribed to particular associations (Becker et al. 2020). HP3’s internal taxonomy is also harmonized with two mammal trees (Bininda-Emonds et al. 2007, Fritz et al. 2009), facilitating analyses that seek to account for host phylogenetic structure while testing hypotheses about viral ecology and evolution (e.g. Becker et al., Farrell et al., Olival et al. 2017, Washburne et al. 2018, Guth et al. 2019, Park 2019, Albery et al. 2020, Mollentze and Streicker 2020). HP3 was also validated against GMPD 1.0.

#### Shaw

Recent work by Shaw *et al*. built a host-pathogen edgelist by combining a systematic literature search with cross-validation from several of the above-mentioned datasets (Shaw et al. 2020). Similar to the construction of HP3, the authors started with lists of known pathogenic bacteria and viruses found in humans and animals. They then conducted Google Scholar searches pairing pathogen names with disease-related keywords, followed by manual review of search results. For well-studied pathogens they limited their manual review to a subset of the top 200 most “relevant” publications as determined by Google. From the resulting literature searches, the authors compiled 12,212 interactions between 2,656 vertebrate host species (including, but not limited to, mammals) and 2,595 viruses and bacteria. GMPD2, EID2, and the Global Infectious Diseases and Epidemiology Network (GIDEON) Guide to Medically Important Bacteria (Gideon Informatics, Inc. and Berger 2020) were used to validate the host-pathogen associations. The dataset is available as a static flat file through figshare and the project GitHub repository (Shaw et al. 2020). Host-pathogen associations are provided alongside pathogen metadata (e.g., genome size, bacterial traits, transmission mode, zoonotic status) and diagnostic method (i.e., PCR, pathogen isolation, pathology). The dataset also includes a comprehensive host phylogeny, developed specifically for the study using nine mitochondrial genes for downstream analyses of host phylogenetic similarity and host breadth.

### A reconciled mammal virome dataset

Though some of these datasets were validated against each other during production, they are sometimes used for cross-validation in analytical work (Albery et al. 2020), and some studies have generated a study-specific *ad hoc* reconciled dataset (Farrell et al. 2020, Gibb et al. 2020), no work has been published with the primary aim of reconciling them as correctly, comprehensively, and reproducibly as possible. Dynamic datasets like EID2, and recent datasets like Shaw, can inherently draw on a greater cumulative body of scientific work. This could mean they include most of the data captured by previous efforts, yet we found there are substantial differences among all four datasets. In isolation, we expect that these differences could impact ecological and evolutionary inference in ways that are difficult to quantify, with special relevance to significance thresholds in hypothesis-testing research (i.e., different datasets may confer different power to statistical tests). In unison, we expect that these data could be standardized into one shared format, allowing them to cover a greater percentage of the global virome, a greater diversity of host species, and obviating the need for researchers to either choose between them or implement *ad hoc* solutions that merge them prior to analysis.

To illustrate the potential for comprehensive data reconciliation, we harmonized the four major datasets described here, creating a new synthetic ‘CLOVER’ dataset out of the four “leaves” (which we have made available with this study). To do so, we first harmonized the host taxonomy of all four datasets using the R package ‘taxize’ (Chamberlain and Szöcs 2013), then manually resolved remaining discrepancies. Finally, using the Julia package ‘NCBITaxonomy.jl’ (Poisot 2020), we standardized host and virus taxonomy against the taxonomic hierarchy (Schoch et al. 2020) used as a reference by the National Center for Biotechnology Information’s Taxonomy database (ncbi.nlm.nih.gov). With all four datasets taxonomically consistent, we were able to show that each only covered a portion of the known global mammal virome, even for the most studied hosts and viruses (Figure 1). Our taxonomic harmonization helped reconcile some discrepancies, increasing overlap among the datasets (Figure 2), but notable differences remained. This could confound inference: for example, using a simple linear model, we found that **data provenance** (see Glossary) explained 8.8% of variation in host species’ viral diversity (but only 4.7% after harmonization). When studies report different findings based on slight variation around a significance threshold, readers should therefore wonder whether subtle differences in the underlying datasets might account for such variation.

**Figure 1.**
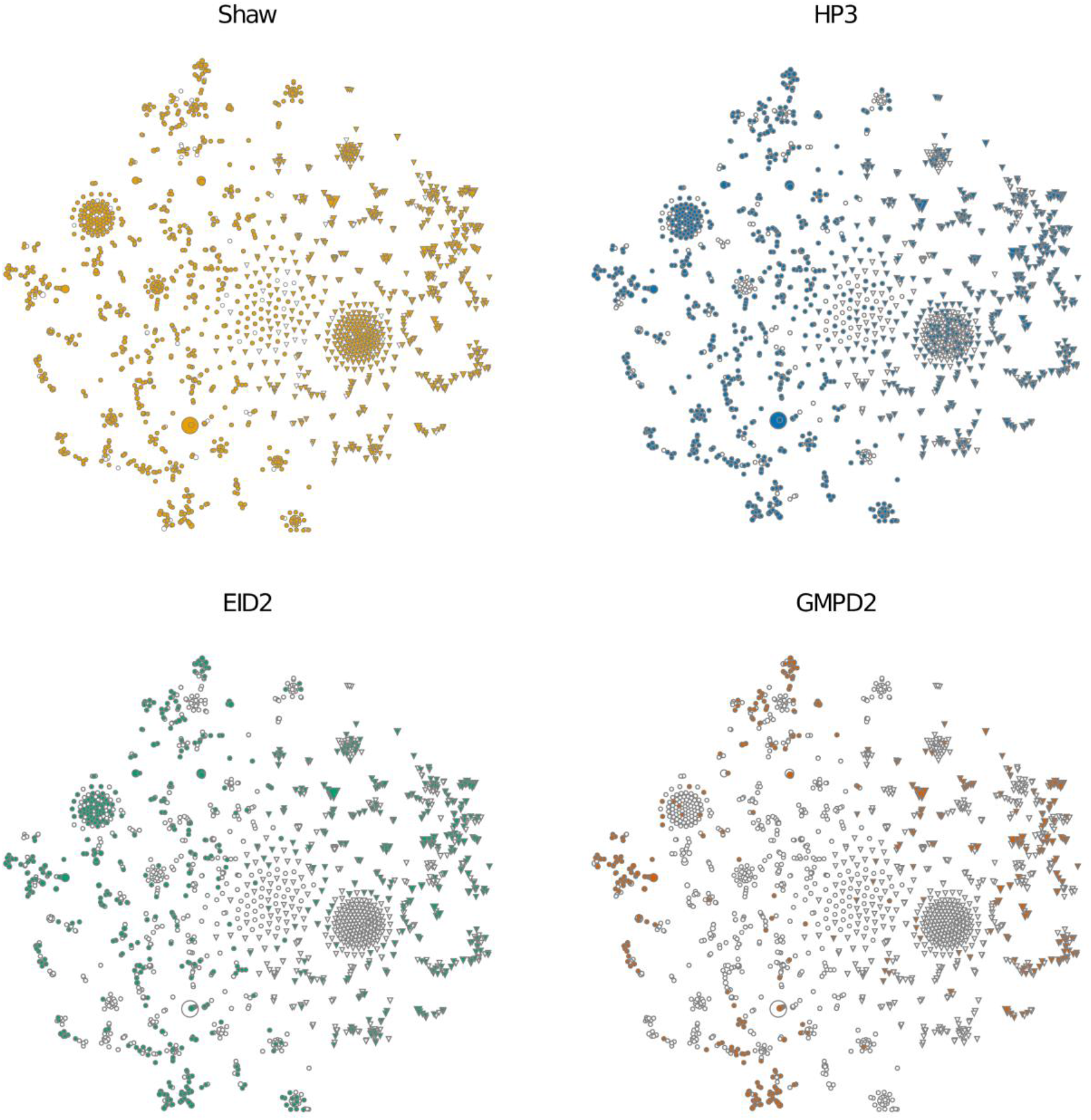
Network representation of the CLOVER dataset. The nodes of the entire CLOVER network have been projected to a two-dimensional space using t-SNE; in each panel, only the nodes found in the dataset are shown in colour. In each dataset, a non-trivial proportion of associations are completely unique and unrecorded elsewhere, even after taxonomic reconciliation. This was the case for 203 of 1360 associations in EID2 (14.9%); 614/2783 in HP3 (22.1%); 269/893 in GMPD2 (30.1%); and 1705/4207 in Shaw (40.5%).

**Figure 2.**
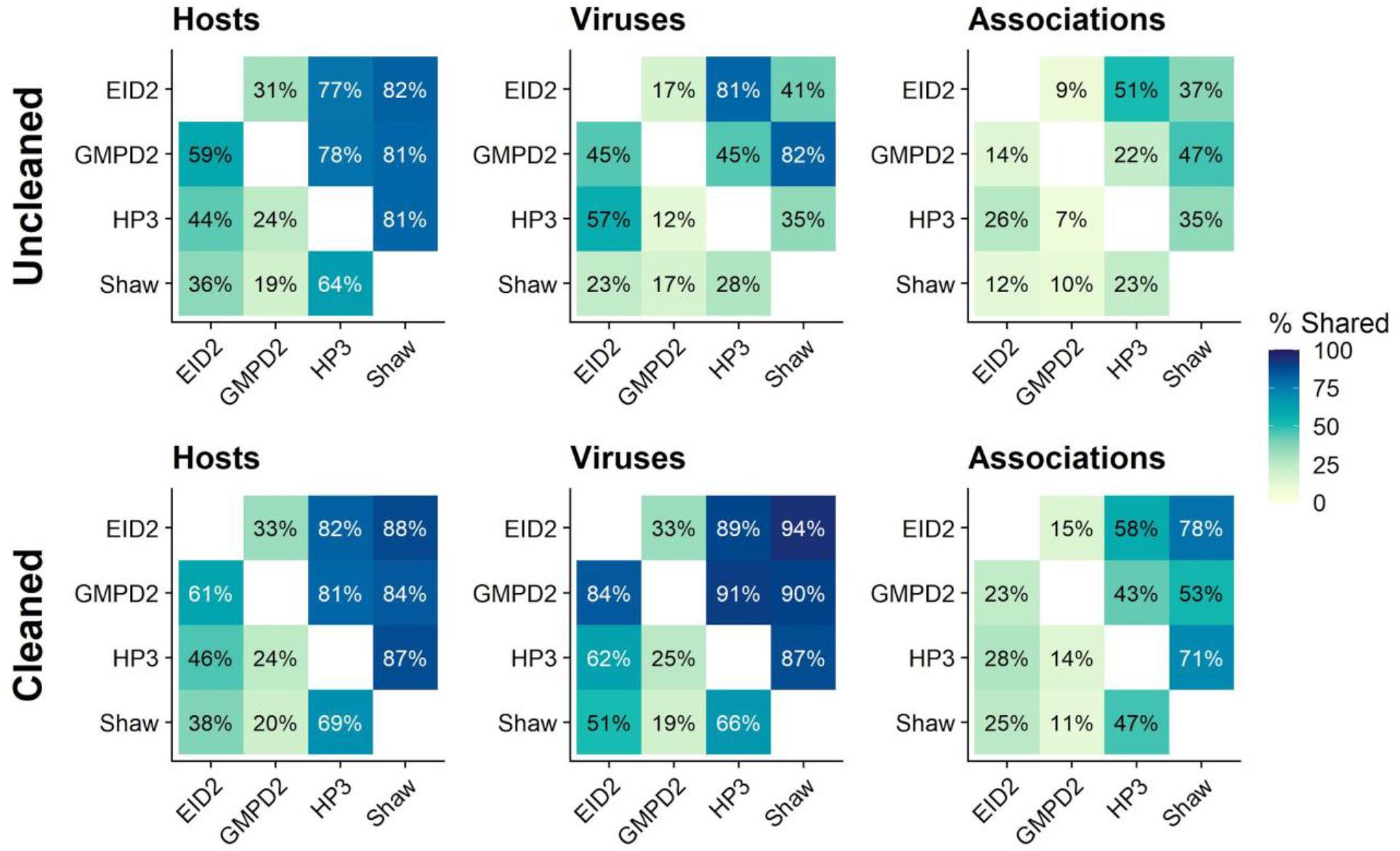
Proportional overlap before and after host taxonomic updating. The percentages and fill colours in these tiles can be interpreted as “% y axis was contained in x axis”; for example, 32% of uncleaned EID2 hosts were also represented in GMPD2, while 47% of cleaned Shaw associations were also contained in HP3. Darker colours represent greater overlap.

Integrated datasets move us a step closer to resolving this uncertainty. The CLOVER dataset covers 1,081 mammal host species and 829 associated viruses. This only represents 16.9% of extant mammals (Burgin et al. 2018) and at most 2.1% of their viruses (Carlson et al. 2019) - perhaps a marginal improvement over the 954 mammal hosts (14.9%) and 733 viruses (1.8%) in the reconciled Shaw sub-dataset, but an improvement nonetheless. The biggest functional gain is not in the *breadth* of the reconciled data, but in its *depth*: the Shaw database records 4,209 interactions among these host and virus species, while CLOVER captures 5,494. Given that previous studies have estimated that 20-40% of host-parasite links are unknown (in GMPD2 (Dallas et al. 2017)), this 30% improvement is notable and shows the value of data synthesis: both building out *and* filling in synthetic datasets will significantly improve the performance of statistical models, which are usually heavily confounded by matrix sparsity (Becker et al., Dallas et al. 2017).

In addition, harmonization of metadata on virus detection methods across datasets enables a greater scrutiny of the strength of evidence in support of each host-virus association. We applied a simplified detection method classification scheme (either serology, PCR/sequencing, isolation/observation, or method unknown) based on descriptions in the source databases or, where these are not provided, adopting the most conservative definition given data source (i.e., EID2 entries derived from NCBI Nucleotide are classified under PCR/sequencing, though they might also qualify for the next strongest level of isolation/observation; whereas entries derived from PubMed are classified under method unknown). Of the 5,494 unique host-virus pairs in CLOVER, a total of 2,156 (39%) have been demonstrated using either viral isolation or direct observation and 1,895 (34%) via PCR or sequencing-based methods (with some overlap, as some associations have been reported with both of the above methods). Notably, a substantial proportion (2,257; 41%) are based solely on serological evidence which, although an indicator of past exposure, does not necessarily reflect host competence (i.e. effectiveness at transmitting a pathogen; Gilbert et al. 2013, Lachish and Murray 2018, Becker et al. 2020). These harmonized definitions facilitate investigation of inferential stability using various types of evidence, as well as enabling a best practice of subsetting data for a particular research purpose. For example, serological assays are a much weaker form of evidence if the aim of a study is zoonotic reservoir host prediction, whereas isolation data open new avenues for testing hypotheses about reservoir competence (Becker et al. 2020).

Data synthesis inherently relies on a scientific community that generates new, often conflicting, data. The generation of truly novel data or finding ways to resolve existing observations that are in conflict are two equally viable paths to scientific progress. However, in the current funding landscape, researchers may have a significant incentive to position themselves as creating an entirely “novel” dataset from scratch, even if it partially replicates available data sources, or to focus their limited resources on datasets that improve the depth of knowledge within a narrow scope (e.g., a focus on specific taxonomic groups). But when testing microbiological or eco-evolutionary hypotheses, rather than simply using each newly-published dataset as a benchmark for which one is “most up-to-date,” we suggest a necessary shift in scientific cultural norms towards using synthetic, reconciled data like CLOVER as an analytical best practice. To make this possible, at least a handful of researchers will need to continue the task of stepwise integration, using datasets that synthesize existing knowledge across teams, institutions, and funding programs to fill in critical data with even more detail. The required tasks (e.g., identifying relevant source data, cleaning taxonomic information, harmonizing metadata on diagnostic information or spatiotemporal structure) can be time-consuming but are relatively straightforward to conduct, and can increasingly be automated thanks to the rapid growth of new data and tools for reproducible research (Boettiger et al. 2015, Lowndes et al. 2017, Colella et al. 2020). There is a clear need, and no obvious technical barrier, to invest more effort in data harmonization: engaging in this process as a form of open science will accelerate progress for the entire research community.

### Relevance to future efforts

Here, we showed that a simple data synthesis effort can create a dramatically more comprehensive dataset of mammal-virus associations. However, this is a temporary solution and one that will become less sustainable if similar datasets continue to proliferate or if newer iterations of existing datasets are released, each absorbing different parts of existing efforts. Over the longer term, given global investments in viral discovery from wildlife, static datasets will quickly become out-of-date, and their relation to the most recent empirical knowledge will be left unclear. For example, the CLOVER dataset becomes significantly sparser after 2010, both in terms of the overall number of reported host-virus associations, and the reporting of novel (i.e. previously undetected) associations (Figure 3). This sparseness is most likely due to time lags between host-virus sampling in the field, the reporting or publication of associations, and their eventual inclusion in one of the component datasets, and suggests that CLOVER may now be missing up to a decade’s worth of known host-virus data. In the near term, microbiologists and data scientists may want to approach the task of data reconciliation with a much broader scope, and develop a more sustainable data platform.

**Figure 3.**
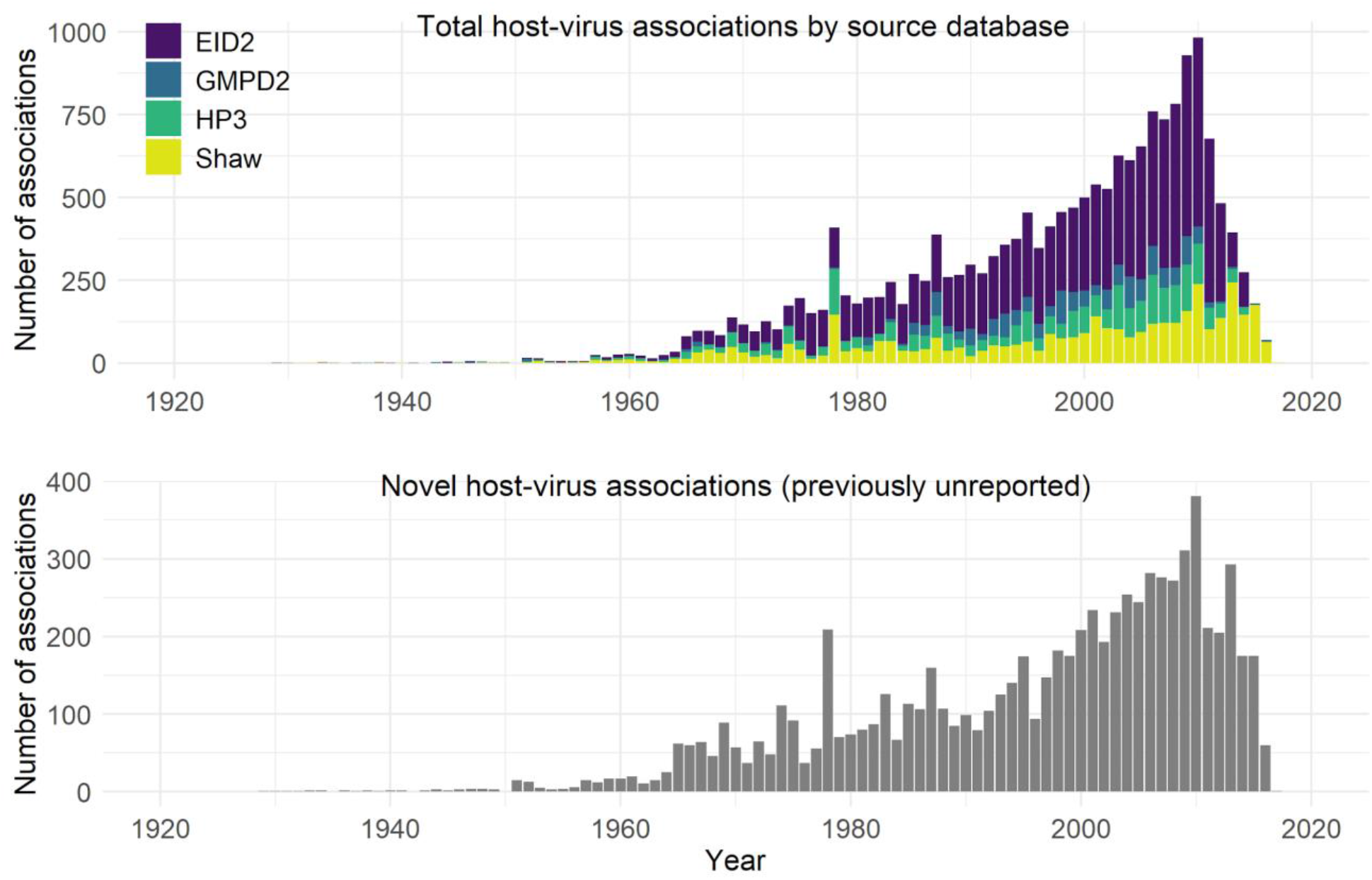
Temporal trends in reporting of host-virus associations in the CLOVER dataset. Bar graphs show, for each year, the total number of reported associations coloured by source database (which can include duplicates of the same association reported over multiple years; top graph) and the number of novel unique associations (i.e. previously unreported; bottom graph). Years reflect the date when an association was reported, either in a published paper or report (for literature-based records) or to the NCBI Nucleotide database (EID2 only).

Scaling up the aggregation of host-virus association data will not be easy, but is not an insurmountable endeavour. We suggest working backwards from the intended end product: the goals outlined here are best served by a central system (with an online access point to the consumable data), spanning the information available from multiple data sources (which demands backend engines drawing from existing databases, while tracking data provenance and ensuring proper attribution). Further, the most valuable data resource would be easily updatable by practitioners (which demands a portal for manual user input or an Integrated Publishing Toolkit to work from flat files). For users, these data should be accessible in a programmatic way (i.e., through a web API allowing for bulk download and/or other interfaces like an R package), help analysts build reproducibility (through versioning of the entire database, or of a specific user query), and offer predictable formats (through a data specification standard devised by a multidisciplinary group).

Fortunately, the field of ecoinformatics has the capacity to help inform this design and development process. Massive bioinformatic data portals like the Global Biodiversity Informatics Facility (gbif.org), the Encyclopedia of Life (eol.org), and the Ocean Biodiversity Information System (obis.org) all offer most of the functionalities we outline here, though they are aimed at slightly different forms of biodiversity data. More recent contributions dedicated to ecological network data include Global Biotic Interactions (Poelen et al. 2014) (GLOBI, which consumes flat files and formats them), helminthR (Dallas 2016), and mangal (Poisot et al. 2016) (which stores a metadata-rich representation of species interaction networks), all of which reconcile their taxonomy with other databases through the use of unique taxon keys. In short, researchers interested in the global virome need not divert their attention, resources, and effort away from the pressing tasks related to monitoring viral pathogens, but they can leverage existing products, expertise, and capacity in neighbouring fields to bolster their ability to do so. Given the eagerness ecologists have shown to participate in SARS-CoV-2 research, we anticipate that our field may be especially well-poised to jump into this task post-pandemic. We aim, in our current efforts, to lay that groundwork.

An integrated platform for the deposition, curation, archival, and sharing of host-virus associations in a *prêt-à-manger*, metadata-rich format has inherent value for the entire scientific community. When the format of a dataset is well established, it allows for the development of tools that mine the data in real-time. For example, the field of biodiversity studies has adopted the concept of Essential Biodiversity Variables, which can be updated when the underlying data change (Pereira et al. 2013, Fernández et al. 2019, Jetz et al. 2019). Having the ability to revisit predictions about the host-virus network could improve models that assess zoonotic potential of wildlife viruses (Farrell et al. 2020, Mollentze et al. 2020), generate priority targets for wildlife reservoir sampling (Becker et al., Babayan et al. 2018, Plowright et al. 2019), and help benchmark model performance related to these tasks. Beyond training and validation, link prediction models built on these reconciled databases may be used to target future literature searches, shifting from systematic literature searches to a model based approach to database updating. Increased collaboration between data collectors, data managers, and data scientists that leads to better data standardization and reconciliation is the only way to productively synthesize our knowledge of the global virome.

## Acknowledgements

This work was supported by funding to the Viral Emergence Research Initiative (VERENA) consortium, including NSF BII 2021909 and a grant from Institut de Valorisation des Données (IVADO). The authors thank Noam Ross, Maya Wardeh, and many others for formative conversations about these datasets and for their tireless work making those data available to the research community.

## Data and code availability

The four raw datasets and harmonized CLOVER dataset can be obtained from the archived project repository: https://dx.doi.org/10.5281/zenodo.4435128. Code used to generate the analyses and figures in this study can be found at https://github.com/viralemergence/reconciliation.

## Box 1.

**Glossary**.

*Association data:* a format that records ecological interactions between a host and symbiont (an *association*) in the form of an edgelist.

*Data provenance:* The primary literature origin of a particular record or set of records in a synthetic dataset.

*Data reconciliation:* the task of harmonizing the language of a given dataset’s fields and metadata to allow a researcher to merge data of different provenance, and generate a new synthetic product.

*Edgelist:* a table, spreadsheet, or matrix of “links” in a host-symbiont network, where each row records the known association of a different host-symbiont pair.

*Flat file:* a static document in Excel or similar spreadsheet or data format, with no dynamic component (no updating) and all data available from a single file rather than a queryable interface.

*Metadata:* additional data describing focal data of interest and that is relevant to interpretation and analysis. Important examples for host-virus associations include sampling method (for example, serological assay, PCR or pathology), date and geographical location of sampling, and standardized information on host and virus taxonomy.

*Open data:* data that is directly and freely accessible for reuse and exploration without impediment, gatekeeping, or cost restriction.

